# Evolutionary history of Alzheimer Disease causing protein family Presenilins with pathological implications

**DOI:** 10.1101/394270

**Authors:** Ammad Aslam Khan, Bushra Mirza, Hashim Ali Raja

## Abstract

Presenilin proteins are type II transmembrane proteins. They make the catalytic component of Gamma secretase, a multiportion transmembrane protease. Amyloid protein, Notch and beta catenin are among more than 90 substrates of Presenilins. Mutations in Presenilins lead to defects in proteolytic cleavage of its substrate resulting in some of the most devastating pathological conditions including Alzheimer disease (AD), developmental disorders and cancer. In addition to catalytic roles, Presenilin protein is also shown to be involved in many non-catalytic roles i.e. calcium homeostasis, regulation of autophagy and protein trafficking etc. These proteolytic proteins are highly conserved, present in almost all the major eukaryotic groups. Studies on wide variety of organisms ranging from human to unicellular dictyostelium have shown the important catalytic and non-catalytic roles of Presenilins. In the current research project, we aimed to elucidate the phylogenetic history of Presenilins. We showed that Presenilins are the most ancient of the Gamma secretase proteins and might have their origin in last common eukaryotic ancestor (LCEA). We also demonstrated that these proteins have been evolving under strong purifying selection. Through evolutionary trace analysis, we showed that Presenilin protein sites which undergoes mutations in Familial Alzheimer Disease are highly conserved in metazoans. Finally, we discussed the evolutionary, physiological and pathological implication of our findings and proposed that evolutionary profile of Presenilins supports the loss of function hypothesis of AD pathogenesis.

## 1. Introduction

Presenilin is a multipass transmembrane protein with nine transmembrane domains (Laudon et al., 2005; Spasic et al., 2006). Presenilin proteins are the major catalytic component of Gamma secretase, a multi-complex protease involved in the catalysis of many membrane proteins (De Strooper et al., 1998; Zhang et al., 2000). Presenilin performs this catalytic activity with the aid of other three Gamma Secretase proteins namely Nicastrin, APH1 and PEN2 (Goutte et al. 2002; Edbauer et al. 2003; Takasugi et al. 2003). In human, there are two Presenilin paralogs, Precesnilin-1 and Presenilin-2 showing 67% sequence similarity (Levy-Lahad et al., 1995). Both these Presenilin proteins are ubiquitously expressed in many human tissues including brain, heart, kidney and muscle (Lee et al., 1996). Inside the cells, Presenilin is abundantly found in Endoplasmic reticulum and Golgi bodies (Area-Gomez et al., 2009; De Strooper et al., 1997; Walter et al., 1996). An important aspect of Presenilin is that under physiological conditions, it undergoes endoproteolysis generating N and C terminal fragments (NTF and CTF respectively) (Brunkan et al., 2005; Thinakaran et al., 1996) involving amino acids 292-299 (Fukumori et al., 2010; Podlisny et al., 1997). This endoproteolysis event is considered important for the Gamma secretase activity. However, there are some Presenilin mutants in which though endoproteolysis is defective but they still exhibit enzymatic activity (Jacobsen et al., 1999; Steiner et al., 1999). There are more than 90 substrates of Gamma Secretase/Presenilin of which APP and the Notch receptor are the predominant substrates (Beel and Sanders, 2008; McCarthy et al., 2009). It is the defect in Presenilin based proteolysis in APP that leads to AD because of the over production of the neurotoxic Aβ42 peptide, which triggers inflammatory responses in brain leading to neuronal dysfunction and ultimately cell death (Duff et al. 1996; Jacobsen et al. 1999; Zhang et al. 2011). However, it is just recently that validity of the amyloid hypothesis as the sole explanation of this AD is questioned and an alternative explanation have been proposed according to which mutations in Presenilin proteins cause loss of many important Presenilin functions (loss of function mutations) in the brain leading to neurodegeneration and dementia, the hallmark features of AD (Kelleher and Shen, 2017; Shen and Kelleher, 2007; Sun et al., 2017).

Quite recently, it has been shown in various studies that in addition to acting as a major protease involved in the regulated intramembrane proteolysis, Presenilin is involved in many non-catalytic processes too. For instance, the role of Presenilin in calcium homoeostasis has been well documented. Presenilin mutations are connected with the defects in calcium signaling in both neuronal and non-neuronal cells and more importantly, also in FAD patients with Presenilin mutations (LaFerla, 2002; Popugaeva and Bezprozvanny, 2013; Supnet and Bezprozvanny, 2010). Presenilin is also involved in the regulation of autophagy in Gammas secretase independent way. The mutations in Presenilin have been shown to be involved in the accumulation of immature autophagic vesicles. The defects in autophagy due to Presenilin mutations contribute towards the neurodegeneration in AD patients because of excess neuronal death (Menzies et al., 2015; Nixon and Yang, 2011; Ohta et al., 2010). Finally, Presenilin have also been found to be involved in the trafficking of proteins in both Gamm secretase dependent (Cai et al., 2003; R. Wang et al., 2006) and independent way (Scheper et al., 2004; Suga et al., 2004).

Though it is known that Presenilins are present in vast variety of eukaryotes, very little work have been done to explore the evolutionary history of this important protein. Even the studies that have been conducted in this direction, are limited either to elucidate the functional aspects of Presenilins in evolutionarily diverse variety of model organisms, or are done before the advent of next generation sequencing technology and hence their coverage is quite limited (Murray et al. 2000; J. Wang et al. 2006; Otto et al. 2016). In this regard, ours is the first such study, elucidating the phylogeny and evolutionary history of Presenilins by exploring their origin and evolution and discussing the physiological implication of the evolutionary profile they exhibits. In our surge, we found that Presenilin is a very ancient protein and might be present in LCA of eukaryotes. We also showed that Presenilin have been evolving under strong purifying selective pressure. Finally we hypothesized that evolutionary history of Presenilin does not support the amyloid hypothesis as an explanation of the etiology of Alzheimer dieses.

## 2. Materials and Methods

### 2.1 Identification and Extraction of Gamma secretase proteins

For the extraction of protein sequences of four Gamma secretase proteins, Presenilin, Nicastrin, APHI and PEN-2, BLAST search tool was used (Altschul et al., 1990). Orthologs of each of these four proteins in human genome were used as query sequence to find the corresponding ortholog and paralog in total 47 species, including each major eukaryotic group, from primates to unicellular protists (Supplementary file 1). For all those proteins with not a well-defined annotation, the result was confirmed with reverse BLAST (Wall et al., 2003) and confirming the putative domain structure from CDD database (Marchler-Bauer et al., 2015). For those species whose genome has been sequenced but still not incorporated in NCBI database or is partially incorporated, BLAST search was conducted from the individual genome database websites of each such species.

### 2.2 Phylogenetic analysis

Phylogenetic analyses were performed using Mega6 (Tamura et al., 2013). Briefly, a total of 153 protein sequences were selected from 47 different species ranging from primates to simplest of protists. Proteins with partial sequences were not included in the analysis. The sequences were aligned by using ClustalW tool employed in MEGA software with affine gap penalties of 5 for gap opening and 1 for gap extension (Chenna et al., 2003). The evolutionary history was inferred using the Neighbor-Joining method (Saitou and Nei, 1987). The analysis gave rise to an optimal tree with the sum of branch length = 39.25074069 (see the figure legend for a detailed account). For rate of selection analysis, coding sequences of Vertebrate Presenilin-1and Presenilin-2 gene were extracted from NCBI genome database (Supplementary file 3). Different approaches were adopted to find the rate of evolution of vertebrate Presenilin proteins. Pairwise comparison of the number of synonymous nucleotide substitutions per synonymous (dS) site and non-synonymous nucleotide substitutions per non-synonymous site (dN) was carried out by using Nei-Gojobori method (Nei and Gojobori, 1986). In addition to pairwise methods, the dN/dS ratio in different branches of the maximum-likelihood tree was estimated using the codon-based genetic algorithm implemented in the GA-BRANCH program available at the Datamonkey server (http://www.datamonkey.org/help/GABranch.php). This approach assigns each branch to an incrementally estimated class of dN/dS ratios without requiring a specification of the branches *a priori* (Kosakovsky Pond and Frost, 2005a). For selection analysis for each Codon, sliding window approach was incorporated in SLAC method employed at the Datamonkey server (Kosakovsky Pond and Frost, 2005b, 2005c). Evolutionary distance between all possible pairs of vertebrate Presenilin paralogs was estimated by Tajima’s relative test employed in Mega6 software (Tajima, 1993; Tamura et al., 2013). Each pair of paralogs was compared with Saccoglossus kowalevskii Presenilin protein sequences which was taken as outgroup.

### 2.3 Evolutionary Trace Analysis (ETA)

For determining the evolutionary conserved residues, evolutionary trace analysis were employed. ETA is a powerful technique that rank the individual amino acids on the bases of their conservation profile. Lower the rank, more conserve the residue and vice versa, with 1 being the minimum rank value (Lichtarge et al., 1996). The ETA results were incorporated for evolutionary structural analysis by employing pyETV and CHIMERA software tools (Lua and Lichtarge, 2010; Pettersen et al., 2004). The heatmaps for the for the figurative representation of ETA results are generated by using conditional formatting tool embedded in Microsoft excel (Meyer and Avery, 2009).

## 3. Results

### 3.1 Distribution and extent of duplication of Gamma secretase proteins across eukaryotes

In our surge to elucidate the evolutionary history of metazoan Presenilins, we started with investigating the distribution of four Gamma secretase proteins, Presenilin, APH-1, PEN-2 and Nicastrin in 46 species representing all the major groups in eukaryotes (Supplementary file 1 and 2). BLAST analysis revealed that these proteins though are distributed in diverse array of eukaryotic the, the extent of distribution is not uniform (Fig.1). For instance, Presenilin seemed to be present in almost all the eukaryotic lineages with the exception of two alveolates (Vitrella brassicaformis and Tetrahymena thermophile) and one excavate (Naegleria gruberi). On the other hand, other gamma secretase proteins included in this study i.e. Nicastrin, Pen-2 and APH-1 were relatively less widely distributed then Presenilin. Except of APH-1 which seems to be absent in plants, all three the three Gamma secretase proteins are present in the three major divisions of eukaryotes e.g. plants, animals and fungi (Fig.1). Among the major eukaryotic groups Alveolates showed least number of Gamma secretase proteins i.e. in all the five species we included in our sample, both APH-1 and Nicastrin were absent while Pen-2 was present in only one Alveolate species e.g in Hammondia hammondi. Similarly in four Excavates we included in our BLAST analysis, Pen-2 is present in only one species i.e. in Trichomonas vaginalis while both APH-1 and Nicastrin are absent in all four. After opisthokonta, plants and ameobazoa are the eukaryotes in which we found pre-dominant presence of Gamma secretase proteins.

**FIG. 1.**
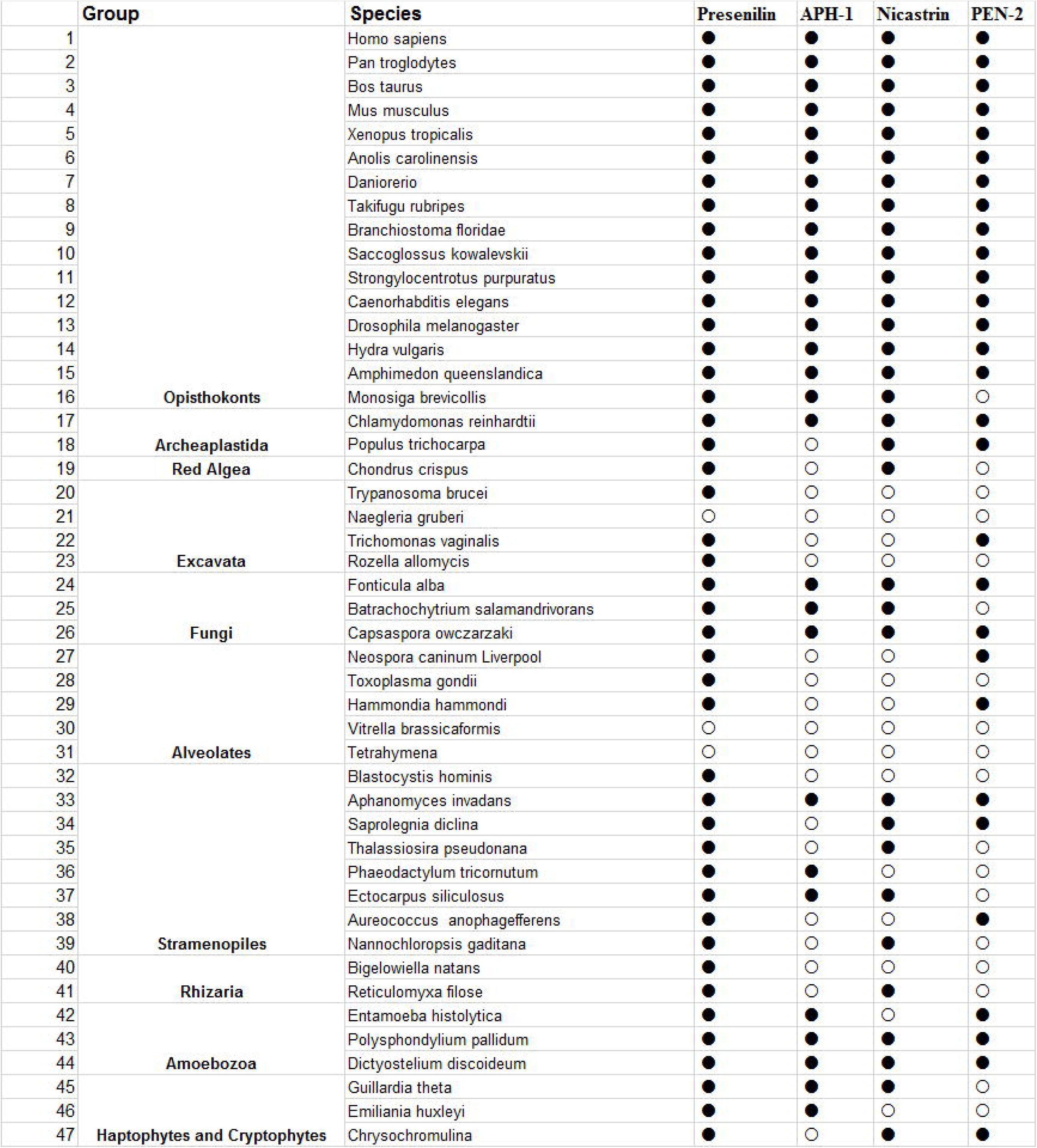
Distribution of Gamma secretase proteins in 47 species belonging to major eukaryotic groups from human to protists. Solid and hollow circles represent the presence and absence of the respective proteins respectively.

In addition to exploring the distribution of Gamma secretase genes, we also investigated the extent of duplications of Presenilins and their other counterpart Gamma secretase genes in eukaryotes. Interestingly, though the number of paralogs for all four genes were different in different species, it seems that these genes underwent less duplications in general. In vertebrates for instance, neither Pen-2 nor Nicastrin seem to have their counterpart paralog with the only exception of Xenopus in which these proteins had one counterpart paralog each. Similarly, both Presenilin and APH-1 are represented by two paralogs in vertebrates e.g. Presenilin-1 and Presenilin-2 and APH-1A and APH-1B. Only in mice, APH-1 have a third paralog e.g. APH-1C. In invertebrate metazoans, all four proteins are in unduplicated form (singleton) with the exception of C.elegans in which Presenilin are represented by three paralogs (Sel-12, Hop-1 and Spe-4). Similar to the case of multicellular organisms, we did not see significant gene duplication based expansion of three of the Gamma secretase proteins (Nicastrin, APH1 and Pen-2) in major unicellular groups. However Presenilin did expanded to some extent (with maximum of four paralogs) in some of these organisms (Fig 2 and Supplementary file 4).

**FIG. 2.**
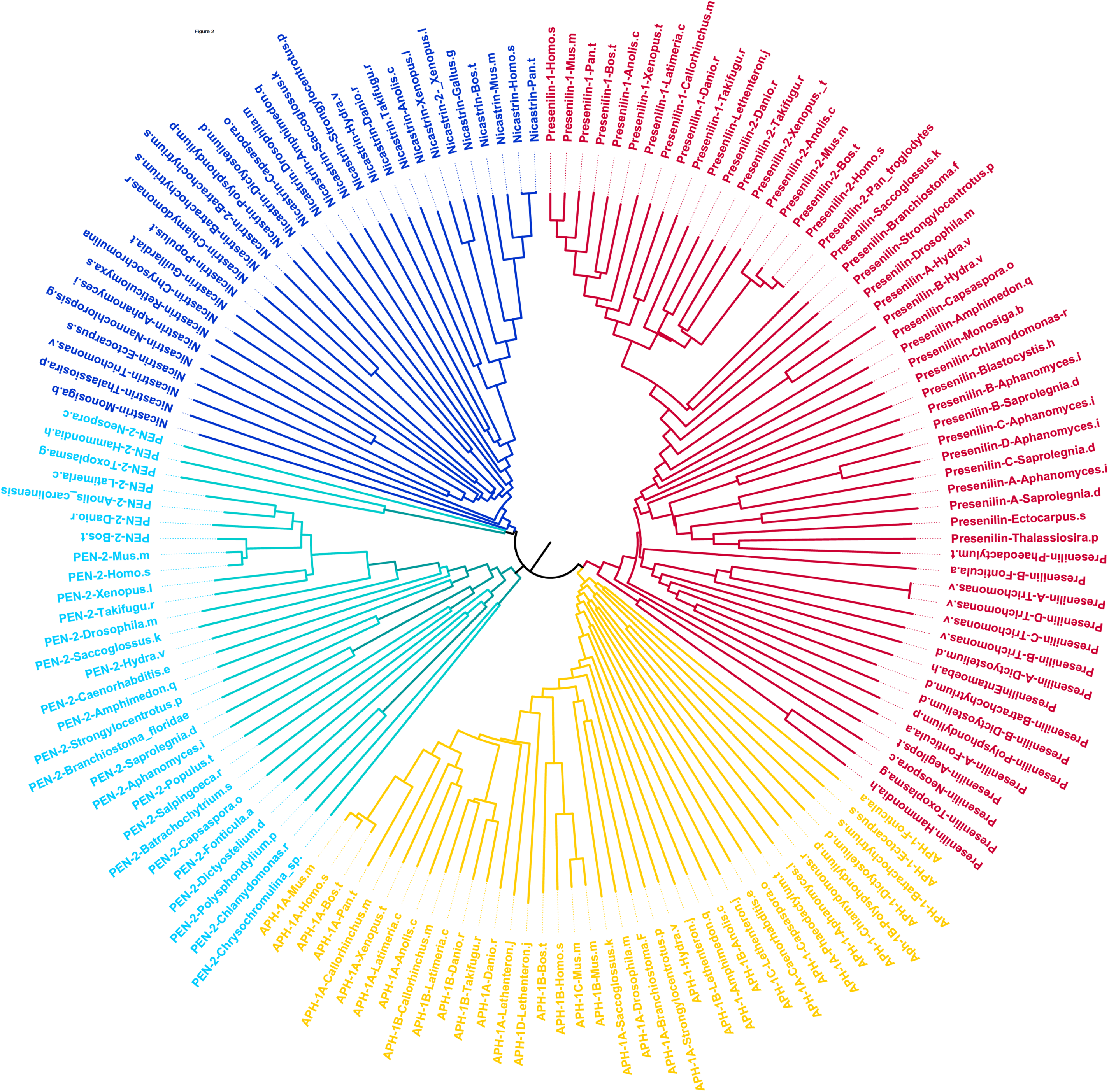
Evolutionary Tree of Gamma Secretase proteins. Phylogenetic tree depicting the phylogenetic history of four Gamma secretase proteins. The evolutionary history was inferred using the Neighbor-Joining method (Saitou and Nei, 1987). The optimal tree with the sum of branch length = 39.25074069 is shown. The tree is drawn to scale, with branch lengths in the same units as those of the evolutionary distances used to infer the phylogenetic tree. The evolutionary distances were computed using the p-distance method (Nei and Kumar, 2000) and are in the units of the number of amino acid differences per site. The analysis involved 153 amino acid sequences. All ambiguous positions were removed for each sequence pair. There were a total of 2897 positions in the final dataset. Evolutionary analyses were conducted in MEGA 6 (Tamura et al., 2013).

### 3.2 Rate of evolution of Presenilins

Exploring the rate of evolution of proteins is an important part towards the elucidation of their evolutionary history. To this direction, we attempted to investigate the rate of evolution of Presenilins, the most widely distributed Gamma secretase protein in eukaryotes. We opted for different approaches including determining the net dn/ds difference in Presenilins, branch site based method (GA-branch) to estimate the lineage specific rate of evolution and sliding window method to estimate the rate of evolution of each site along the whole length of Presenilin proteins (fig. 3 A-D and supplementary file 5). Interestingly, in all these selection analysis, we witnessed both Presenilin predominantly evolving under purifying selection. Finally, we employed Tajima’s relative rate test to find out if there is any significant difference between rates of evolution of two Presenilin proteins (fig. 3 E). In this test, we compared the extent of substitutions in both vertebrate Presenilin paralogs by taking the protein sequence of a third species as outgroup in which the respective protein exists as unduplicated singleton e.g. the invertebrate Saccoglossus kowalevskii. The results of Tajima’s test showed that there is no significant difference in evolutionary rate between two paralogs.

**FIG. 3.**
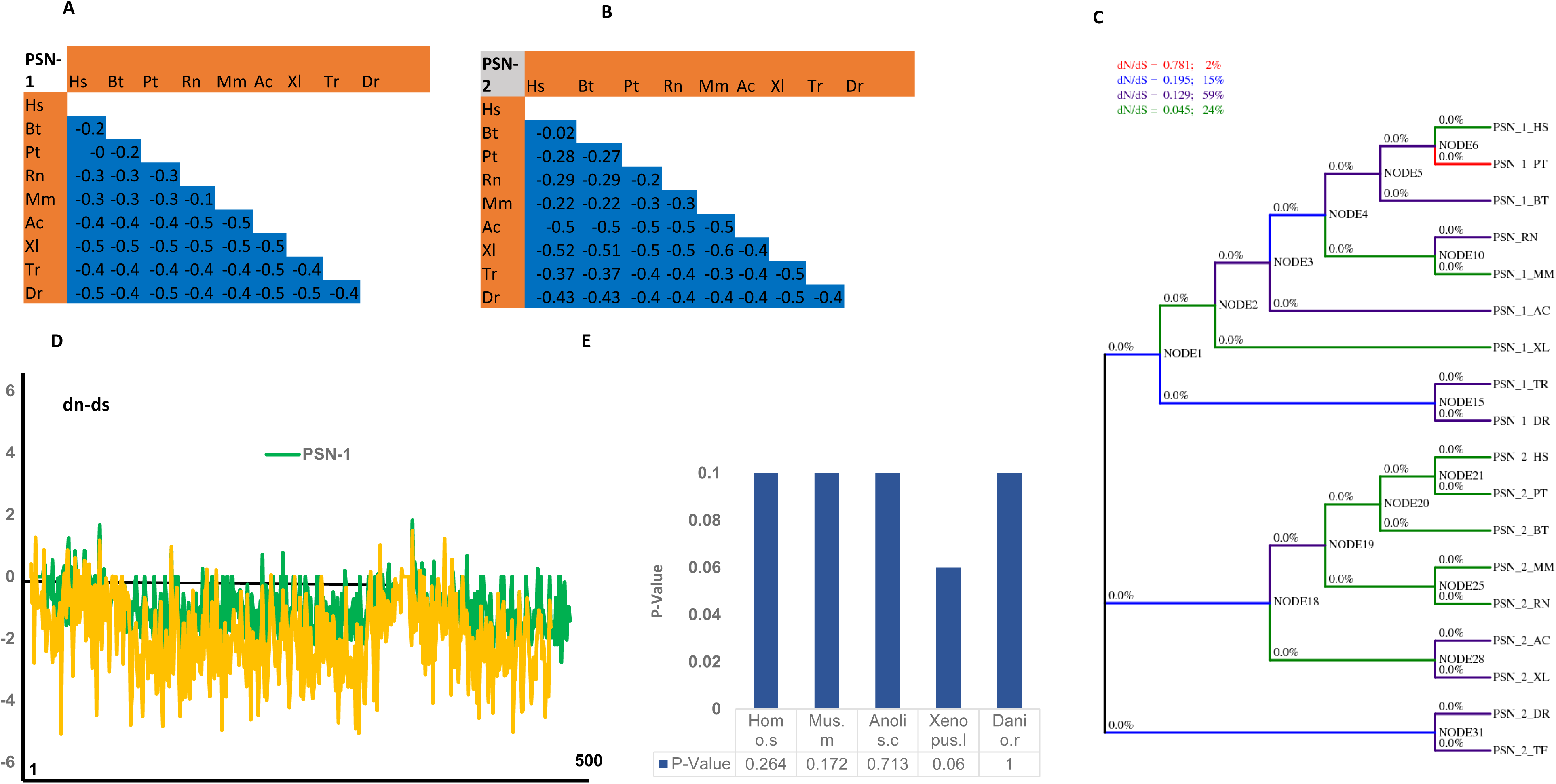
Selection analysis of Presenilin proteins in Vertebrates. (A and B) For both Presenilin-1 and Presenelin-2, the difference between the nonsynonymous and synonymous differences per site between different species are shown. Analyses were conducted using the modified Nei-Gojobori (assumed transition/transversion bias = 2) model. A negative value means the sequence is under purifying selection. (C) GA-Branch based analysis of lineage specific selective pressure in Presenilin genes. A cladogram is shown with maximum-likelihood estimates of lineage specific *d*N/*d*S during vertebrate Presenilin evolution. Percentages for branch classes in the legend reflect the proportion of total tree length (measured in expected substitutions per site per unit time) evolving under the corresponding value of *dN*/*dS* with letters A, B, C, D and E represents each branch class. (D) Sliding window display of nonsynonymous and synonymous differences for each site (codon) in vertebrate Presenilins. SLAC method from datamonkey server is employed for this analysis. (E) Tajima’s relative rate test to depict the difference in rate of evolution of two Presenilin paralogs in vertebrates with Presenilin protein sequence of Saccoglossus kowalevskii as outgroup.

### 3.3 Evolutionary Traces in metazoan Presenilins

Evolutionary Trace Analysis (ETA) is a powerful phylogenetic tool to identify evolutionarily conserved residues in protein sequences. The main principle of ETA is that mutations occurring in distant branches in a phylogenetic tree are more important than those occurring at closely related branches. In this way, ETA not only contribute towards our understanding of the evolutionary history of a protein but also provide a mean to identify biologically important residues (Lichtarge et al., 1996). So, in order to identify the highly conserved regions and residues in metazoan Presenilin, we employed ETA. With the help of ETA analysis, we identified 103 residues in Presenilins which were conserved (with rank value of 1) throughout metazoans from primates to choanoflagellates (fig. 4A and supplementary file 6). We also looked for the location of these significantly ranked residues along the whole length of Presenilin protein. Interestingly, majority of the highly ranked residues were present in the transmembrane part of the protein (fig. 4B and C). In order to see the biological importance of these traces, we looked for the site directed mutation analysis by making use of Uniprot (Pundir et al., 2017) (fig. 4D). As was expected, a significant number of mutating sites (60%) were highly conserved among the metazoans as depicted by their very low rank values rank vale of 1 or 2.

**FIG. 4.**
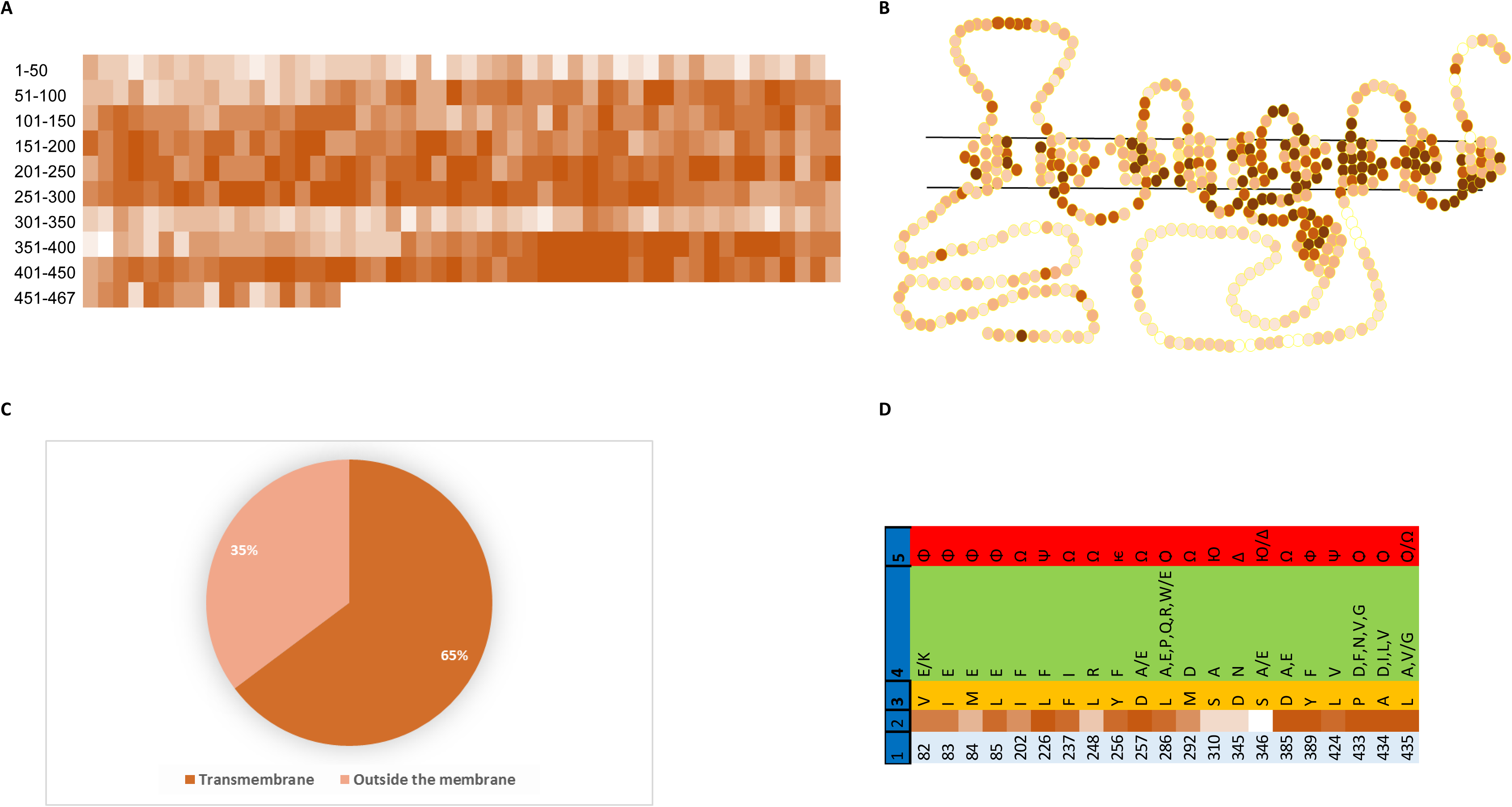
Evolutionary traces analysis of Presenilin-1 protein**. (A)** A heatmap of evolutionary traces found in metazoan Presenilin. The color shades from dark brown to white correspond to most conserved to least conserved residues. The TA is carried out in by using 15 species, covering major metazoan groups from primates to choanoflagellates.(B) A Schematic heatmap based representation of Presenilin-1 protein across the cell membrane. The color shades corresponds to same conservation profile as mentioned in A and B. The scheme of arrangement of Presenilin across the membrane is taken from the work of Zhang et al. (Zhang et al., 2013). (C) Pi graph showing the distribution of highly conserved residues –as depicted by the ETA rank profile-in the transmembrane and non-transmembrane part of the Presenilin protein. (D) List of site directed mutations which have been carried out in Preselin-1 gene. Heatmap shown in column 2 depicts the conservation profile of mutated sites with color shades from dark brown to white correspond to most conserved to least conserved residues. Phenotypic effects due to these mutations are given in columns 5. Φ= Loss of interaction with GFAP, Ω= Abolished protease activity, Ψ= increased protease activity, □= alters gamma secretase specificity, □= Reduced Notch processing, Ю= Abolished PKA signaling. Δ= Abolished caspase cleavage.

### 3.4 Alzheimer Disease and Presenilin Evolution

Mutations in Presenilin gene are among the hallmark causes of Alzheimer disease. In our surge for elucidation evolutionary history of Presenilin, we were keen to investigate its implication in the pathogenesis of Alzheimer disease. To this direction, we made use of the uniprot database for natural variants (Pundir et al., 2017) in human Presenilin-1 protein causing familial Alzheimer disease (FAD). When we applied the ETA analysis on these AD/FAD causing sites, we were amazed to see a predominant number of sites exhibited very high level of conservation (70%) throughout metazoans from human to choanoflagellates (fig. 5A and B). Of these 70% of rank 1 residues, majority lie in the alpha helix as we observed by the structural analysis in which we mapped the ETA based ranked residues on the existing human Presenilin-1 structure (fig. 5C). Similarly, the surface view of the structure reveals the predominant presence of these conserved sites in the hydrophobic region of the protein which is in consistent with our analysis that a big majority of these residues lie in the transmembrane part of the protein. (fig. 5D).

**FIG. 5.**
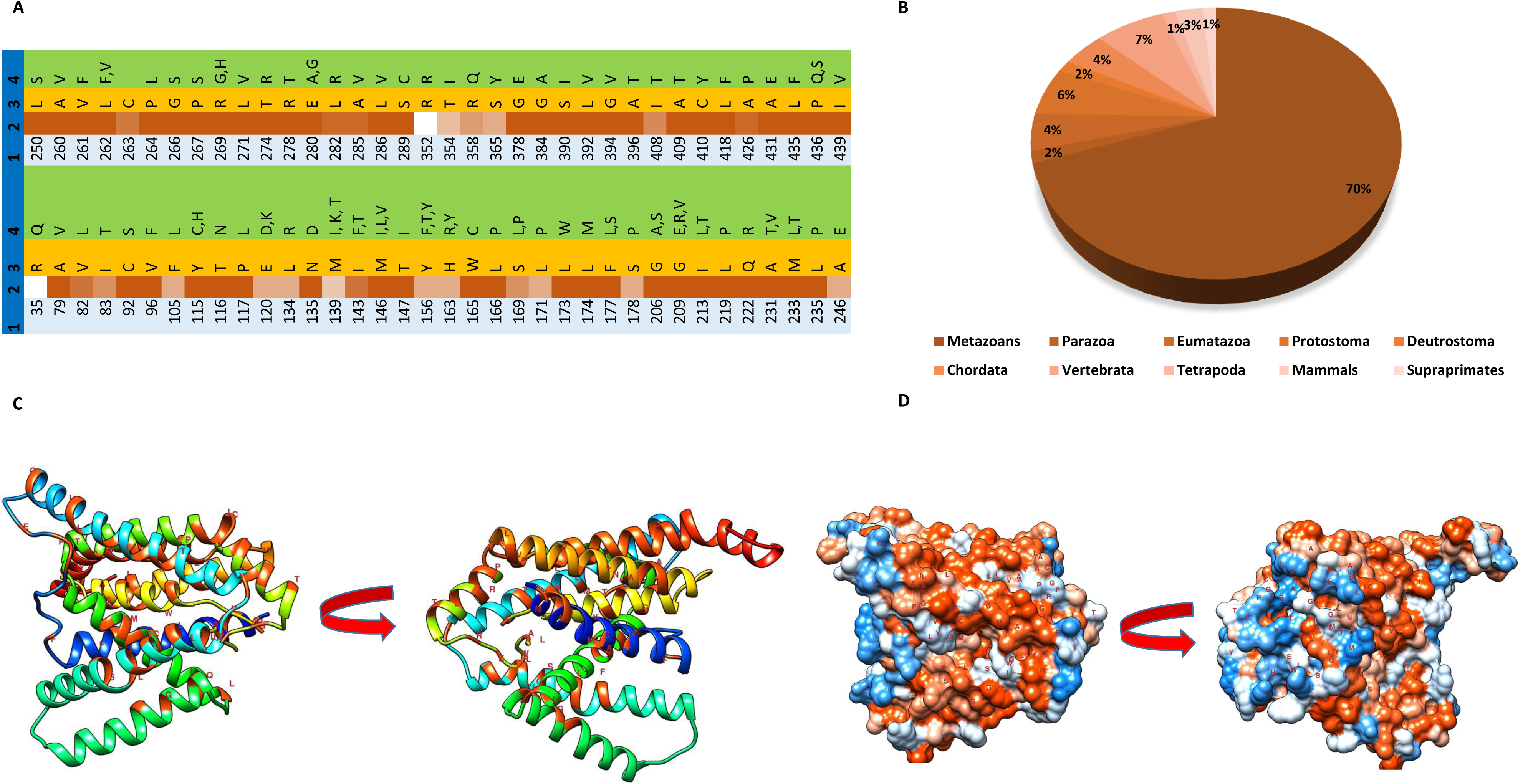
Evolution of Presenilin and Alzheimer disease. (A) List of Presenilin-1 mutations leading to AD along with conservation profile of each mutated site in the form of heatmap. The color shades from dark brown to white correspond to most conserved to least conserved residues. (B) Pi graph showing the extent of conservation of evolutionary traces mutation in whom leads to AD. (C and D) ETA based structural analysis of Presenilin protein by using Cryo-EM structure of human Presenilin-1 (5NF2) as template. Both ribbon (C) and surface (D) cartoons are shown with the red spots depicting the location of highly conserved residues (traces) susceptible for AD. ETA based ranks are generated by comparing protein sequences of 15 species, covering major metazoan groups from primates to choanoflagellates.

## 4. Discussion

Gama secretase is a protein complex that along with β-Secretase is involved in the successive proteolysis of amyloid precursor protein producing amyloid β-protein whose abnormal deposition in the brain leads to AD. Of the four proteins making γ-Secretase, it is the catalytic role of Presenilin which is most prominently involved in the AD pathogenesis. Because of the important role Presenilin play in AD and its other catalytic and non-catalytic roles, it performs in evolutionarily diverse eukaryotes, we considered it important to explore the evolutionary history of this protein and its implication in the roles it performs. A protein BLAST based search for all the γ-Secretase proteins showed that these proteins are widely distributed in all the eukaryotic lineages with Presenilin showing the widest distribution, reflecting its vital role. Among all the species we included in our sample for protein evolution study, only one excavate (Naegleria gruberi) and Ciliates were the groups lacking Presenilin gene in their genome. It is understandable for parasitic species like Naegleria to lack such genes as parasitic organisms tend to have reduced genome because of repeated gene loss during the course of evolution (JACKSON, 2015; Sakharkar and Chow, 2005; Slamovits, 2013). But absence of Presenilin in Tetrahymena (a free living ciliate) may be surprising though. However, Ciliates seem to lack other important substrates of Presenilin too i.e. Amyloid precursor protein and Notch (data not shown). This reflects the possibility of coevolution of Presenilin with its substrates though more study is needed to explore this aspect of Presenilin evolution. Whether the absence Presenilin in Ciliates is result of gene loss or a representative of a feature inherited from a simpler common ancestor cannot be determined at the moment because of the unresolved phylogeny of the basal protist groups (Baldauf, 2003; Keeling et al., 2005; Parfrey et al., 2006). Interestingly, we did not find a single species in which any of the other three γ-Secretase proteins was present when Presenilin was absent. It not only means that Presenilin is the most important protein as for as activity of γ-Secretase is concerned but also mean that perhaps in the eukaryotic LCA, γ-Secretase activity was exhibited solely by Presenilins. It is interesting to notice that Amyoloid Beta protein though is highly conserved in metazoans, it is not present in vast number of unicellular organisms. On the other hand, in the same way as is Presenilin, Notch proteins are present throughout the eukaryotes, in metazoans, protists, plants and fungi. Therefore we can say that in LCEA, the main proteolytic function of Presenilins was the catalysis of Notch related proteins while the catalysis of amyloid beta is rather a metazoan innovation. Likewise, incorporation Presenilin to form Gamma secretase complex is also most probably a metazoan innovation, contributing towards the diversity of its proteolytic activity to fulfil the requirements of a complex multicellular organization. The independent evolution of these two catalytic activities of Presenilin is also reflected by the fact that mutations in Presenilin protein that affect the Amyloid beta proteolysis does not primarily affect proteolysis of Notch by Presenilin (Capell et al., 2000; Kulic et al., 2000). The conservation of Asp-257 and Asp-385 throughout the eukaryotic life, from human to hydra to all the protists we included in our analysis, signifies the pivotal role of Presenilins as a protease. This preservation of catalytic residues in species in which Presenilin is the only representative protein of Gamma secretase confirms the fact that assembly of Presenilin to form Gamma secretase is not required for performing its proteolytic activity (Beher et al., 2001). The conservation of Aspartic acid residues in those species in which Presenilin is the only of the Gamma Secretase protein mean that in LCEA, a proto-Presenilin could perform its catalytic activity independent of the Gamma secretase activity. Likewise, as Presenilin perform its non-catalytic functions i.e. protein trafficking, calcium homeostasis and apostasies etc. independent of Gamma secretase activity (Capell et al., 2000; Otto et al., 2016), presence of Presenilin in these simple organisms also hints towards the ancient origin of non-catalytic role of Presenilin which it performs independent of Gamma secretase. So a protein which is present such a diverse group of organisms and performing a variety of important functions of both catalytic as well as non-catalytic nature, must also be evolving under some considerable functional constrains. And this is what we observed in our evolutionary rate analysis too. We employed different strategies to estimate the rate of evolution of vertebrate Presenilin i.e. estimation of net dn/ds difference in Presenilins, Hyphy based estimation of the lineage specific rate of evolution and sliding window method to estimate the rate of evolution of each site along the whole length of Presenilin proteins. As expected, all these strategies proved that Presenilin protein have been evolving under strong purifying selection. This was further confirmed by the Tajima’s relative rate test which showed that even duplication of vertebrate Presenilin did not affect this trend and both the Presenilin paralogs went on evolving under negative selection pressure. This is also in accordance with the general understanding that the older genes (and hence present in wide range of organisms) and the genes more important for the survival of the organism evolve slowly than the younger ones (Albà and Castresana, 2005; Domazet-Loso and Tautz, 2003; Hirsh and Fraser, 2001; Wilson et al., 1977; Wolf et al., 2009).

One of the important functional aspects of Presenilin is the structure-function relationship of Presenilin (Bai et al., 2015). The study of the topology of Presenilin has revealed important aspects of this structure function relationship. For instance, studies conducted towards this direction have shown the importance of transmembrane portions of Presenilin involved in the catalytic activity of Gamma secretase. For instance, studies based upon substituted cysteine accessibility method (SCAM) and Nuclear Magnetic Resonance (NMR) have shown the importance of TMD1, TMD5, TMD6, TMD7 and TMD9 in the catalysis of transmembrane proteins (Sato et al., 2008, 2006, Tolia et al., 2008, 2006; Watanabe et al., 2010). In this direction, we also looked to see if we can see any selective constrain also on transmembrane domains of Presenilin as compared to the cytosolic or luminal portions. As was expected, the ETA analysis also showed that a significant majority (65%) of the highly conserved amino acids (with ETA rank value = 1) were present in the transmembrane region of the protein. This conservation of the transmembrane portion of the Presenilin reflects the functional constrain TMDs exhibit by providing a hydrophilic catalytic pore in the hydrophobic milieu for an efficient catalysis of the membrane proteins. Intrigued by the evolutionary history of Presenilin, we were curious to look at the evolutionary profile of amino acid sites in Presenilin whose mutation give rise to Familial Alzheimer disease (FAD). This evolutionary analysis of FAD causing mutations gave some interesting results. For instance, as was expected from structure-function evolutionary analysis, majority of these conserved FAD causing sites were located in the alpha helical transmembrane part of the Presenilin. In addition, many of these mutations seem to be exposed to the hydrophilic aqueous environment as depicted by the surface view of structure analysis. Intriguingly, of the total amino acid sites susceptible for FAD, a significant majority were highly conserved among metazoans from human to choanoflagellates (70%). The genes/loci that are involved in old age diseases usually tend to evolve under lesser functional constrain. It reflects that the sites whose mutation lead to FAD might be involved in performing some highly important functions (Drenos and Kirkwood, 2010; Kirkwood and Austad, 2000). These functions may involve eliciting the signaling cascades and networks controlling physiological manifestations as complex as learning and memory. In other words, we can say that the mutation in these sites might have disturbed important physiological functions in addition to disruption of normal amyloid catalysis. In addition it has been shown that genes involved in a disease due to dominant negative mutations evolve under purifying selection (Blekhman et al., 2008). Interestingly, it has been proposed recently that only amyloid hypothesis cannot account for the occurrence and progression of AD. According this alternative hypothesis, mutations in Presenilin leading to AD/FAD should rather be considered as loss of function dominant negative mutations rather than gain of function mutations as significant number of these mutations severely impair the functioning of Gamma secretase instead of only increasing Aβ42 production (Kelleher and Shen, 2017; Sun et al., 2017; Watanabe and Shen, 2017). This later hypothesis is in conformity with our findings. Therefore, the outcome of our evolutionary analysis showing the ancient origin of Presenilins, their evolution under strong negative selection and strong conservation of the amino acid sites involved in AD pathogenesis favors idea that through loss of function mutations, Presenilins play a dominant negative role leading to AD pathogenesis. Our findings also favor to explore candidates other than amyloid beta also to elucidate the role of Presenilin in AD etiology not only to gain more mechanistic details of AD pathogenicity but also to formulate better therapeutic strategies for this devastating disease.

## 5. Conclusion

Presenilins are type II membrane proteins involved in the proteolysis of quite an important number of proteins. Of the four Gamma secretase proteins, Presenilins are the most primitive one. Their primitive origin means that they may be present in the LCEA and may be the only representative of Gamma secretase activity in LCEA, performing both catalytic as well as non-catalytic roles. Presenilin have been evolving under strong purifying selection representing the physiological vitality of this protein. Remarkably, the Presenilin residues susceptible for AD show significantly high conservation in metazoans manifesting the involvement of these residues in important physiological processes. The mutation in such functionally constrained residues would eventually lead to AD like complex and multifaceted disorder. Based upon their evolutionary profile we can predict that involvement of Presenilin in AD pathogenicity cannot be ascribed to a single cause like amyloid beta production but other factors should also be explored for more effective therapeutic solutions.

## Competing interest

We have no competing interest for this article.

## Author’s contribution

AAK conceived the idea. AAK and BM extracted and analyzed the data, AAK wrote the article.

## Funding

This research was done through the support from by Higher Education Commission of Pakistan (HEC) and Barret Hodgson University, Karachi, Pakistan.

